# Glycosylated GM-CSF expands B-1b cells and B-1b plasma cells and programs them for immunosuppression

**DOI:** 10.64898/2026.02.02.703206

**Authors:** Haisam Alattar, Arpa Aintablian, Ina. N. Eckert, Haroon Shaikh, Laura Cyran, Zheng Liu, Bhupesh Prusty, Fabian Imdahl, Elisabeth Zinser, Andreas Beilhack, Nadine Hövelmeyer, Björn E. Clausen, Tom Gräfenhan, Florian Erhard, Manfred B. Lutz

## Abstract

The myeloid growth factor granulocyte-macrophage colony-stimulating factor (GM-CSF) exhibits paradoxical pro- and anti-inflammatory functions, but the factors determining these divergent outcomes remain unclear. Here, we report that this functional divergence is controlled by its glycosylation. Murine recombinant fully glycosylated GM-CSF (rgGM-CSF) specifically induces immunosuppressive cell types, whereas its recombinant non-glycosylated counterpart (rngGM-CSF) promotes effector immune cells. Using single-cell ATAC-sequencing and flow cytometry, we show that rgGM-CSF has a previously unrecognized ability to effectively expand IL-10^+^ LAG-3^+^ PD-L1^+^ B-1b plasma cells (PCs) with immunosuppressive properties and self reactive natural IgM secretion. Although rgGM-CSF also promotes the expansion of hematopoietic stem and progenitor cells (HSPCs) and monocytic myeloid-derived suppressor cells (M-MDSCs), adoptive transfer experiments demonstrate that the rgGM-CSF-induced B-1b PCs are responsible for an IL-10-dependent long-term protection in mice from experimental autoimmune-encephalomyelitis (EAE). Our data suggest that glycosylation enhances the systemic bioavailability and activity of GM-CSF and promotes the expansion of immunoregulatory cells rather than pro-inflammatory myeloid effector cells. Together, these results demonstrate that the ‘dual activity’ of GM-CSF is controlled by its glycosylation, resulting in opposing immune functions. These findings support a re-evaluation of human rgGM-CSF (regramostim) as a potential therapeutic strategy for immunosuppression in transplantation and autoimmune diseases.

**Key points:**

- Glycosylated GM-CSF promotes B-1b cells and B-1b plasma cells expansion and establishes their long-term imprinting as IL-10^+^ LAG3^+^ PD-L1^+^ natural IgM secreting regulatory cells.
- Albumin binding enhances the systemic activity of glycosylated GM-CSF in generating regulatory B-1b plasma cells.
- Glycosylated GM-CSF injections into mice expand M-MDSCs, but their suppressive iNOS production is only maintained short-term.
- Non-glycosylated GM-CSF injections preferentially promote expansion of pro-inflammatory effector monocytes and neutrophils.

**Graphical Abstract:** 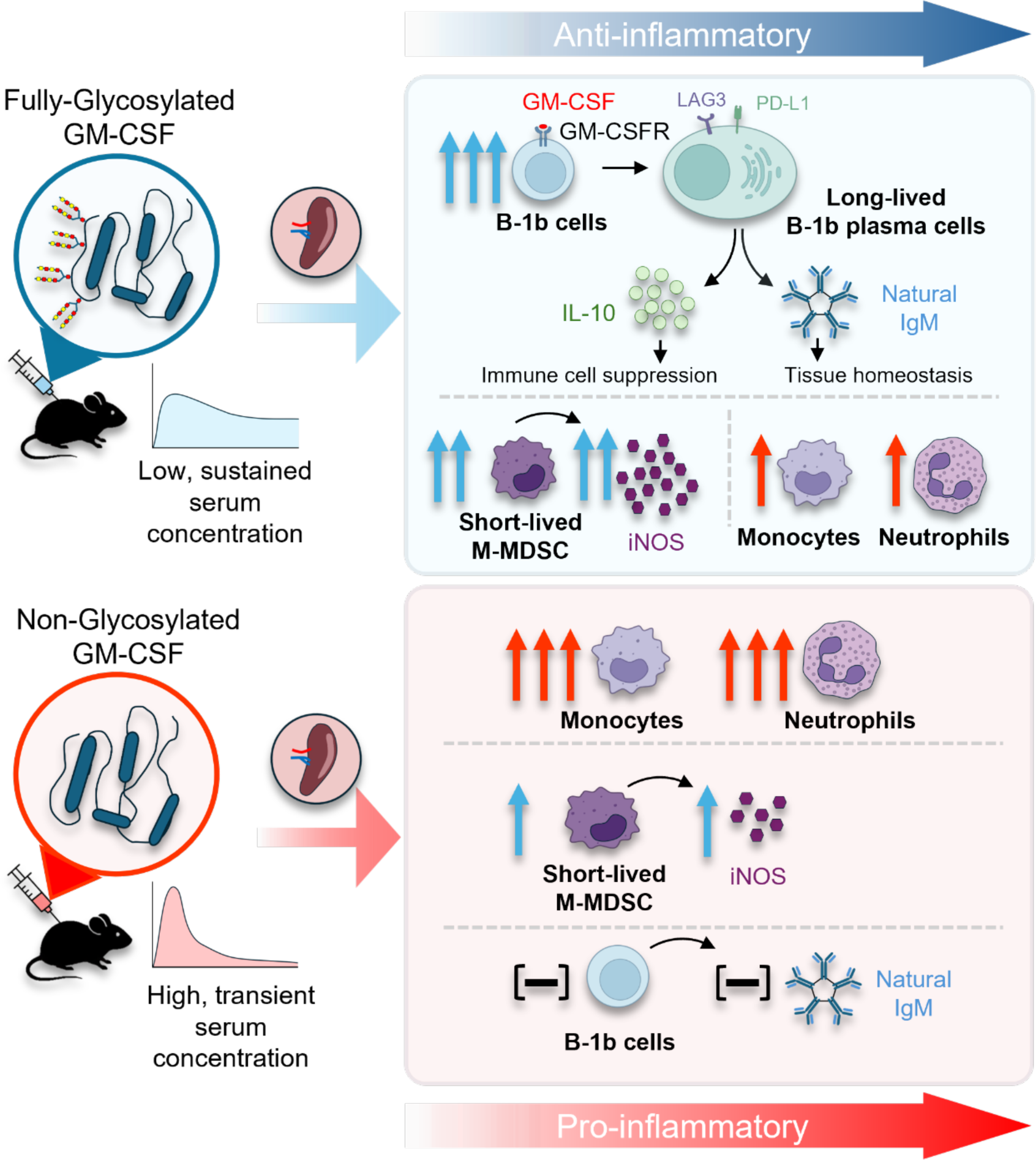

## Introduction

Granulocyte-macrophage colony-stimulating factor (GM-CSF) was initially described as a myeloid growth factor on bone marrow (BM) cells ^1–3^. The partially glycosylated product sargramostim, is approved to promote myelopoiesis in patients after stem cell transplantation ^4,5^. Although, GM-CSF appears dispensable under steady-state conditions, it is a critical pro-inflammatory cytokine during acute inflammations or infections. Consequently, GM-CSF is used as a vaccine adjuvant in tumor patients, exerting effects through neutrophil and monocyte expansion, with monocytes further developing into monocyte-derived dendritic cells (MoDCs) and macrophages ^2,6,7^. The clinical efficacy of GM-CSF variants is dictated by their biochemical properties, as sargramostim and the non-glycosylated molgramostim effectively expanded myeloid effector cells in tumour patients, whereas the fully glycosylated regramostim showed poor efficacy ^8^. These variable outcomes have historically been attributed to dosage or administration route ^9^, yet the underlying molecular mechanisms remain poorly defined.

The functional duality of GM-CSF is particularly evident in chronic disease. Sustained GM-CSF release in cancer patients has led to tumor immune evasion and unfavourable clinical outcomes ^10,11^. Similarly, in chronic inflammatory environments, GM-CSF promotes the accumulation of monocytic and granulocytic myeloid-derived suppressor cells (M-MDSCs, G-MDSCs), which inhibit immune responses via inducible nitric oxide synthase (iNOS), Arginase-1 (Arg-1), and IL-10 ^12–14^. GM-CSF administration has been reported to induce anti-inflammatory responses in mice exhibiting clinical signs of type 1 diabetes or experimental autoimmune thyroiditis ^14–16^ and to reduce Crohn’s Disease in clinical trials ^17^, but again with varied outcomes ^18^. Conversely, GM-CSF neutralization or receptor blockade provides clinical benefits in chronic autoimmunity as multiple sclerosis ^19^ and rheumatoid arthritis ^20,21^. Overall, this ambiguity underscores a fundamental gap in our understanding of how GM-CSF glycoforms dictate pro- or anti-inflammatory outcomes and how various immune subsets respond to these distinct variants.

Recent studies showed that innate-like B cell subsets, such as B-1a cells, respond to autocrine GM-CSF during bacterial infection to secrete polyreactive, pathogen-neutralizing IgM.^22,23^. B-1 cells mainly develop during fetal and neonatal stages and are maintained via self-renewal, with limited replenishment from adult BM.. Two subsets, CD5⁺ B-1a and CD5⁻ B-1b cells, have been identified. Functionally, B-1 cells are the primary source of self-specific natural IgM, which is essential for the clearance of apoptotic debris and tissue homeostasis. Furthermore, B-1 cells can exert immunoregulatory effects under steady-state conditions and during inflammation via IL-10, LAG-3, PD-L1, PD-L2, CD200, and CTLA-4 ^24–27^. Despite these well-established roles, the contribution of GM-CSF to B-1 cell biology remains insufficiently understood.

We previously showed that repeated injections of rgGM-CSF attenuate experimental autoimmune encephalomyelitis (EAE), which was initially linked to higher M-MDSCs levels ^28^. Although adoptive transfer of bulk BM M-MDSCs cultures, generated *in vitro* with GM-CSF, replicated this protection ^28^, we later observed that M-MDSCs disappear within 48 hours post-injection ^29^. Given that these bulk cultures contain hematopoietic stem and progenitor cells (HSPCs) ^30^, we hypothesized that the long-term protective effect observed two weeks post-EAE induction resulted from the transfer of GM-CSF-modified HSPCs that subsequently differentiated into suppressive MDSCs.

Here, we performed single-cell ATAC-sequencing (scATAC-seq) of splenic cells enriched for HSPCs from rgGM-CSF-injected or uninjected mice. The results indicate a strong extramedullary hematopoiesis in rgGM-CSF-injected mice, with the expected expansion of HSPCs and differentiated myeloid cells, including M-MDSCs. Unexpectedly, we found a massive and persistence expansion of IL-10^+^ LAG-3^+^ PD-L1^+^ B-1b plasma cells (PCs) secreting self-specific IgM, phenotypically resembling to a previously described natural regulatory plasma cells (NRPCs) ^26,31^. Adoptive transfer experiments revealed that rgGM-CSF-modified B-1b PCs, rather than modified HSPCs, mediate IL-10-dependent protection against EAE.Interestingly, rgGM-CSF appeared superior at promoting immunosuppressive cells, M-MDSCs and B-1b PCs, while rngGM-CSF preferentially promoted pro-inflammatory myeloid effector cells. Overall, our findings suggest that GM-CSF glycosylation is a critical determinant of its “dual activity” and highlight the translatable therapeutic potential of fully glycosylated human regramostim for targeted immunosuppression in autoimmunity and transplantation.

## Material and Methods

### Mice and GM-CSF treatment

C57BL/6 and IL-10-β-lactamase reporter (ITIB) mice on a C57BL/6 background ^32^, *Nos2*^-/-^ mice (B6.129P2-*Nos2^tm1Lau^*/J), (B6(Cg)-Tyr^c-2J^/J, B6 albino) and luciferase expressing B6.albion mice were bred and maintained under SPF conditions in the animal facilities of the Institute of Hygiene and Microbiology and the Institute of Virology and Immunobiology of the University of Würzburg, and at the Department of Dermatology, University Hospital Erlangen. Both sexes aged 6-12 weeks were used in the experiments. Male CD19-Cre (B6.129P2(C)-*Cd19^tm1(cre)Cgn^*/J) × IL-10^fl/fl^ (B6.129P2(C)-*Il10^tm1Roer^*/MbogJ) and CD19-Cre mice were bread at the University Medical Center of the Johannes Gutenberg-University Mainz. Mice were injected IP for 10 d with 1µg GM-CSF/d using one of the following GM-CSF forms: rgGM-CSF (MCE^®^, Cat. No. HY-P7069), rngGM-CSF (PeproTech^®^, Cat.No.315-03), rgGM-CSF/Bovine Serum Albumin (BSA) (Serva®, Cat. No.11945), rngGM-CSF/BSA, rgGM-CSF/10% normal mouse serum (Sigma-Aldrich^®^, Cat. No.S7273), rgGM-CSF-containing supernatant from murine GM-CSF-transfected myeloma cells producing saturating GM-CSF levels (GM-sup) ^33^, 8 mg BSA, 10% normal mouse serum or phosphate-buffered saline (PBS) as a vehicle control. The animal experiments were performed with permission and under control of the local authorities (Regierung von Unterfranken AZ 54-2532.1-29/08, 54-2532.2-6/08, 55.2-2531.01-67/10, 55.2.2-2532-2-200, and 55.2.2-2532-2-1157).

### EAE induction

The EAE induction protocol was adapted from ^34^. Each mouse was injected SC into the flanks with 100 µl of an emulsion containing 150 µg MOG_35–55_ peptide (MCE^®^, Cat. No. HY-P1240A) in PBS emulsified with with 50 µl of Complete Freund’s Adjuvant (CFA; Sigma-Aldrich^®^, Cat. No. F5881-10ML) containing heat-killed *Mycobacterium tuberculosis* (100 mg/10 ml of CFA; BD Difco^®^, Cat. No. 10218823). At d0 and d2, each mouse was injected IP with 200 ng of pertussis toxin (Enzo^®^, Cat. No.BML-G100). EAE clinical symptoms were scored as follows: 0, no symptoms; 0.5, mild tail weakness; 1, partially limp tail; 2, completely limp tail; 2.5, limp tail with mild hind-limb weakness (wobbling gait); 3, complete hind-limb paralysis; 4, forelimb weakness and rapid breathing; 5, moribund or death.

### Single-cell ATAC-sequencing

To allow two replicates per condition in a single scATAC-seq run, each group, untreated or injected withrgGM-CSF (10x 1µg/d, as GM-sup), included one control mouse (B6(Cg)-Tyr^c-2J^/J, B6 albino) and one firefly luciferase transgenic mouse (B6.luc), both matched for genetic background, age and sex ^35^. At d11, spleen single-cell suspensions of mice from each group were pooled at a 1:1 ratio before enrichment of HSPCs by cell sorting (FACS ARIA III, Becton Dickinson) of Lin^-^ (Ter119, B220, CD4, CD8 and CD11b) c-Kit^+^ cells. The luciferase sequence flanked by the two primers, Luc 1A: 5’ -TCAAAGAGGCGAACTGTGTGTG -3’ and Luc 2B: 5’ - GGTGTTGGAGCAAGATGGAT -3’, was used as a genetic barcode to annotate cells to their respective animals and to bioinformatically analyze individual mice within each group separately. Single-cell ATAC-seq libraries were prepared from sorted HSPCs from untreated and rgGM-CSF-treated mice using the Chromium Next GEM Single Cell ATAC Reagent Kit v1.1(10x Genomics) according to the manufacturer’s protocol (CG000209, Rev D). Briefly, nuclei were isolated from freshly collected sorted HSPCs using the 10x Genomics Nuclei Isolation protocol (CG000124, Rev G) with minor modifications. Nuclei were counted and assessed for integrity using a Countess II FL Automated Cell Counter (ThermoFisher Scientific). Approximately 10,000 nuclei per sample were loaded onto a Chromium Next GEM Chip H for gel bead-in-emulsion (GEM) generation and barcoding on the Chromium Controller (10x Genomics). Within each GEM, transposition was carried out using Tn5 transposase, which fragments accessible chromatin and simultaneously tags DNA with sequencing adapters and cell-specific barcodes. Following transposition and GEM breakage, the barcoded DNA fragments were purified, PCR-amplified to add sample indices, and quantified using a Qubit fluorometer (ThermoFisher Scientific) and Agilent 2100 Bioanalyzer (Agilent Technologies). Libraries were pooled and sequenced on an Illumina NextSeq 2000 system, aiming for a depth of approximately 25,000–50,000 read pairs per nucleus. After assessing sequencing quality with FASTQC, sequence reads were pre-processed with Cell Ranger ATAC 2.1.0 and mapped to the mouse reference GRCm38, with reads mapping to the luciferase transgene used to assign cells to individual mice. Following quality filtering, 10233 cells from rgGM-CSF-treated mice and 4993 cells from untreated mice were retained for downstream analysis. scATAC-seq data are available at GEO under accession number (GSE317457).

### Single-cell ATAC-sequencing analysis

scATAC-seq data processing, integration, and analysis were performed with R (v4.4.2) using Seurat and Signac packages, following the recommended workflows ^36^. To enable direct comparative analysis between the two datasets, the rgGM-CSF-injected and untreated cells dataset, a unified peak set was generated by merging peaks from the two datasets. Each sample was re-quantified by counting fragments overlapping the union peaks ^37^. High-quality cells were retained based on fragment abundance (3,000–25,000 fragments/cell), fraction of reads in peaks (FriP) > 15%, nucleosome signal < 4, transcription start site (TSS) enrichment > 3, blacklist ratio < 0.05. Dimensionality reduction was performed using Term Frequency-Inverse Document Frequency (TF-IDF) normalization followed by Singular Value Decomposition (SVD). Batch effects were corrected using Harmony (LSI dimensions 2–30), followed by unsupervised clustering using the Louvain algorithm and UMAP visualization. Inferred gene activity scores were calculated using the GeneActivity function in Signac R package from chromatin accessibility, log-normalized and used for differential testing to define the enriched genes of each cluster ^36^. Cell-cell communication was inferred using the CellChat R package on gene activity profiles ^38^. Finally, the top 200 upregulated genes in cluster 2 were identified from gene activity values, and pathway enrichment analysis was carried out using GO Enrichment Analysis powered by PANTHER ^39^.

### MACS sorting for CD138^+^ PCs

Peritoneal cells were collected by peritoneal lavage after the 10 d injection period and washed once with PBS. The spleen was harvested and processed to obtain a single-cell suspension ^40^. Peritoneal and splenic cells (5x10^7^ total) were resuspended in autoMACS™ Rinsing Solution (Miltenyi Biotec, Cat. No. 130-091-222; 2mM EDTA, 0.5% BSA, pH 7.5). The suspension was stained with biotinylated antibodies against CD4, CD8, Ly6C, Ly6G, CD11c, and NK1.1 diluted 1:50 for 20 minutes on ice to deplete T cells, myeloid cells, DCs and NK cells. The cells were then washed and resuspended in 280µl autoMACS™ Rinsing Solution, followed by incubation with 20 µl Streptavidin MicroBeads (Miltenyi Biotec; Cat. No. 130-048-102) for an additional 20 minutes on ice. The labelled cells were depleted by passing the suspension through an LD Column (Miltenyi Biotec, Cat. No. 130-042-901), and the negatively selected, PCs-enriched fraction was collected and washed. Then, the enriched cells were resuspended in the autoMACS™ Rinsing Solution and stained with biotinylated anti-CD138 (diluted 1:50) for 20 minutes on ice. After washing, the cells were incubated for 20 minutes with 20µl Streptavidin MicroBeads, and then underwent a second magnetic separation using an LS Column (Miltenyi Biotec; Cat. No. 130-042-401) to isolate the CD138⁺ PCs. After one wash, the positively selected cells were resuspended in RPMI medium containing 10% FCS and 1x10^5^ cells were adoptively transferred intravenously via the lateral tail vein into mice before EAE induction. This procedure yielded a purity greater than 95%.

### *In vitro* B-1 cell culture

A suspension of C57BL/6 spleen cells was prepared as described in ^40^. Then, 3x10^5^ cells were cultured in a 24-well plate in 1 ml of R10 medium (RPMI 1640 supplemented with 10% heat-inactivated FCS, 2 mM L-glutamine, 50 μM β-mercaptoethanol, and 100 U/ml penicillin/streptomycin). Cells were cultured either without any stimulant or with 3.5 µM rgGM-CSF. Additional wells were cultured with 10 µg/ml purified F(ab’)₂ goat anti-mouse IgM (μ-chain) alone (anti-B cell receptor, anti-IgM, Biolegend^®^, Cat. No.157102), with 25 ng/ml IL-4 alone (PeproTech^®^, Cat. No.214-14), anti-IgM + IL-4 or anti-IgM + IL-4 + rgGM-CSF. Cultures were incubated at 37°C for 4 days, and flow cytometric analysis of B-1 cells was performed on day 4.

### *In vitro* BM M-MDSC generation and LPS/IFN-γ stimulation

M-MDSCs were generated from BM of C57BL/6 or *Nos2*^-/-^ mice as described before ^40,41^. Briefly, BM cells were flushed from the tibia and femur of adult mice, washed once with PBS, and resuspended in R10 media. The single-cell BM suspensions (3x10^6^ cells in 10ml R10 in a 10 cm dish) were then cultured in the presence of 10% GM-sup for 3 days at 37°C and 5% CO_2_. For M-MDSC activation, spleen single cell suspensions were cultured in 24-well plates at 10^6^ cells/ml/well in the presence of 100 ng/ml Lipopolysaccharide (LPS, Sigma-Aldrich^®^, Cat. No. L2880) and 100 U/ml IFN-γ (Immunotools^®^, Cat. No. 12343536) for 16 h as described ^13,14,40,42^.

### *Ex vivo* IL-10 stimulation and staining

A protocol adapted from Bouabe et al. ^32^ was used. Briefly, 5x10^6^ peritoneal and spleen cells from 10x rgGM-CSF (± BSA) injected ITIB mice were stimulated in R10 media containing 50 ng/ml phorbol 12-myristate 13-acetate (PMA, Sigma-Aldrich^®^, Cat. No.P1585) and 1 µg/ml Ionomycin (Sigma-Aldrich^®^, Cat. No.I9657) for 6 h at 37°C/5% CO_2_. Cells were then washed, and 1x10^6^ cells were resuspended in R10 media loaded with 1.3 µM CCF-4 solution (ThermoFisher scientific^®^, Cat. No.K1095) and 3.6 µM probenecid (ThermoFisher scientific^®^, Cat. No.KP36400), followed by incubation for 90 minutes at room temperature in the dark. Cells were subsequently washed and stained with fluorochrome-conjugated antibodies for other markers.

### Flow cytometry staining and analysis

To stain surface markers, cells were incubated for 30 minutes at 4°C with the antibody mix in staining buffer (PBS, 0.1% BSA and 50% 2.4G2 hybridoma supernatant to block FcγRII/III). To stain the intracellular markers, the cells were fixed with 2% formaldehyde or with Fix/Perm buffer (eBioscience™ Foxp3/ Transcription Factor Staining Buffer Set, Cat. No.00-5523-00) for 20 minutes at room temperature. After that, the cells were washed and resuspended in the intracellular antibody mix in permeabilization buffer for 1 hour at room temperature. Samples were acquired with the Attune NxT Flow Cytometer (Thermo Fisher Scientific), and FlowJo 10.10 (Tree Star) was used for data analysis. The following directly conjugated anti-mouse antibodies were purchased from BioLegend^®^: Ter119 FITC (TER-119), B220 FITC or BV650 (RA-3-6B2), CD11b FITC, AF647 or AF700 (M1/70), CD11c PE-Cy7 (N418), CD4 FITC (GK1.5), CD8 FITC (53-6.7), NK-1.1 FITC (PK136), Ly6C BV510 (HK1.4), Ly6G FITC or BV650 (1A8), CD117 PE/Cy7 (2B8), CD16/32 PerCP/Cy5.5 (S17011E), CD34 AF700 (RAM34), Sca-1 BV510 (D7), CD115 AF647 (AFS98), CD19 PerCP/Cy5.5 (1D3/CD19), CD138 BV711 (281-2), CD5 APC (53-7.3), CD43 AF700 (S11), CD3 APC/Cy7 (17A2), PD-L1 PE (10F.9G2). Anti-mouse LAG3 PE/Cy7 (eBioC9B7W), iNOS AF488 (CXNFT) and Fixable Viability Dye eFluor 780 (Cat.No. 65-0865-14) were purchased from eBioscience^®^. Anti-mouse GMCSFR alpha PE (698423) was purchased from R&D systems^®^.

### LEGENDplex™ for mouse GM-CSF and IL-6

Serum samples were collected on d11 from mice that had been injected with PBS, BSA, rgGM-CSF, or rngGM-CSF for 10 days. Serum levels of GM-CSF were then determined using LEGENDplex™ Mouse GM-CSF Capture Bead B9 (Cat. No. 740163) and LEGENDplex™ Mouse IL-6 Capture Bead B4 (Cat. No. 740159) according to the manufacturer’s protocol.

### Detection of IgM natural antibodies against self-antigens by ELISA

To detect natural antibodies in mouse serum, we developed a sandwich ELISA based on ^43^. 96-well half-area microplate were coated overnight at 4°C with 50µl of carbonate coating buffer (100mM NaHCO₃, 33.6mM Na₂CO₃, pH 9.5) containing anti-fibronectin 1 (α-FN1) IgM (1:500, Sigma-Aldrich^®^ clone P1H11), anti-phosphorylcholin (α-PC) IgM (1:5,000, Sigma-Aldrich^®^ clone BH8), or anti-malondialdehyde (α-MDA) IgM (1:10,000, Sigma-Aldrich^®^ Cat. No. SAB5202544). For the IgM standard curve, 50µl of 50ng/ml native mouse IgM was added to the first two wells, which were then serially diluted two-fold, including blank wells. The next day, the plates were washed five times with PBST (PBS with 0.05% Tween-20) and blocked with 2% BSA in PBST for 2h at room temperature while shaking. After washing, 50µl of diluted serum samples were added (1:500 for α-FN1 IgM plates; 1:1,000 for α-MDA IgM and α-PC IgM plates) and incubated for 1h at room temperature with shaking. After another wash, 50µl of HRP-conjugated goat-α-mouse IgM (Invitrogen, No. 31440, 1:10,000) was added and shaken for 1h at room temperature. After a final wash, we added 50µl of TMB substrate (BioLegend^®^, Cat. No.421101) for 20 minutes, then 50µl of 2N H2SO4 to stop the reaction. Absorbance was measured at 450 nm using ThermoFisher Scientific Multiskan FC Type 357 microtiter plate photometer, and IgM concentrations were calculated from the standard curve generated with the native mouse IgM.

### Statistics

Data analysis was done using GraphPad Prism 10.6.1 and Microsoft Excel 365. The specific statistical tests used for each experiment are listed in the figure legends. Differences between groups were considered statistically significant at p < 0.05.

## Results

### rgGM-CSF drives extramedullary hematopoiesis in the spleen and expands B-1b PCs

Previously, we found that on one hand, adoptive transfer of rgGM-CSF bulk BM cultures enriched for M-MDSCs protected from EAE, but on the other hand, M-MDSCs rapidly disappeared 48h after their injection ^28,29^. From this, we hypothesised that GM-CSF-imprinted HSPCs, which are still present as minor contaminants in the M-MDSC cultures, may be responsible for EAE protection. To investigate whether rgGM-CSF triggers epigenetic imprinting of HSPCs toward an immunosuppressive M-MDSC fate capable of conferring EAE protection, we performed scATAC-seq on splenic HSPCs sorted after 10x daily administration of rgGM-CSF. Untreated mice served as controls (Fig. 1A). Notably, the rg-GMCSF injections elicited marked splenomegaly compared with the untreated control (Fig. 1B). The scATAC-seq analysis demonstrated a broad expansion of HSPCs (Fig. 1C). GM-CSF preferentially expanded three distinct myeloid progenitor populations: *Flt3*⁺ common myeloid progenitors (CMPs; cluster 6; *Kit⁺ Cd34⁺ Flt3⁺ Fcgr3⁻ Csf1r^lo^ Ms4a3^lo^*), granulocyte-monocyte progenitors (GMPs; cluster 8; *Kit⁺ Cd34⁺ Fcgr3⁺ Csf1r^lo^ Ms4a3⁺ S100a9⁺ Cd177⁺*), and the cMoP/monocytes (cluster 5; *Kit⁺ Cd34⁺ Fcgr3⁺ Csf1r^hi^ Ms4a3^lo^ S100a9⁻ Ly6c2⁺*), consistent with early monocyte and neutrophil commitment. This HSPCs expansion was confirmed at the protein level by flow cytometry ^44^, and showed expansion of long-term hematopoietic stem cells (LT-HSCs), short-term HSC (ST-HSCs), besides CMPs, GMPs, and cMoPs compared with untreated controls (Suppl. Fig. 1E, F).

**Fig. 1.**
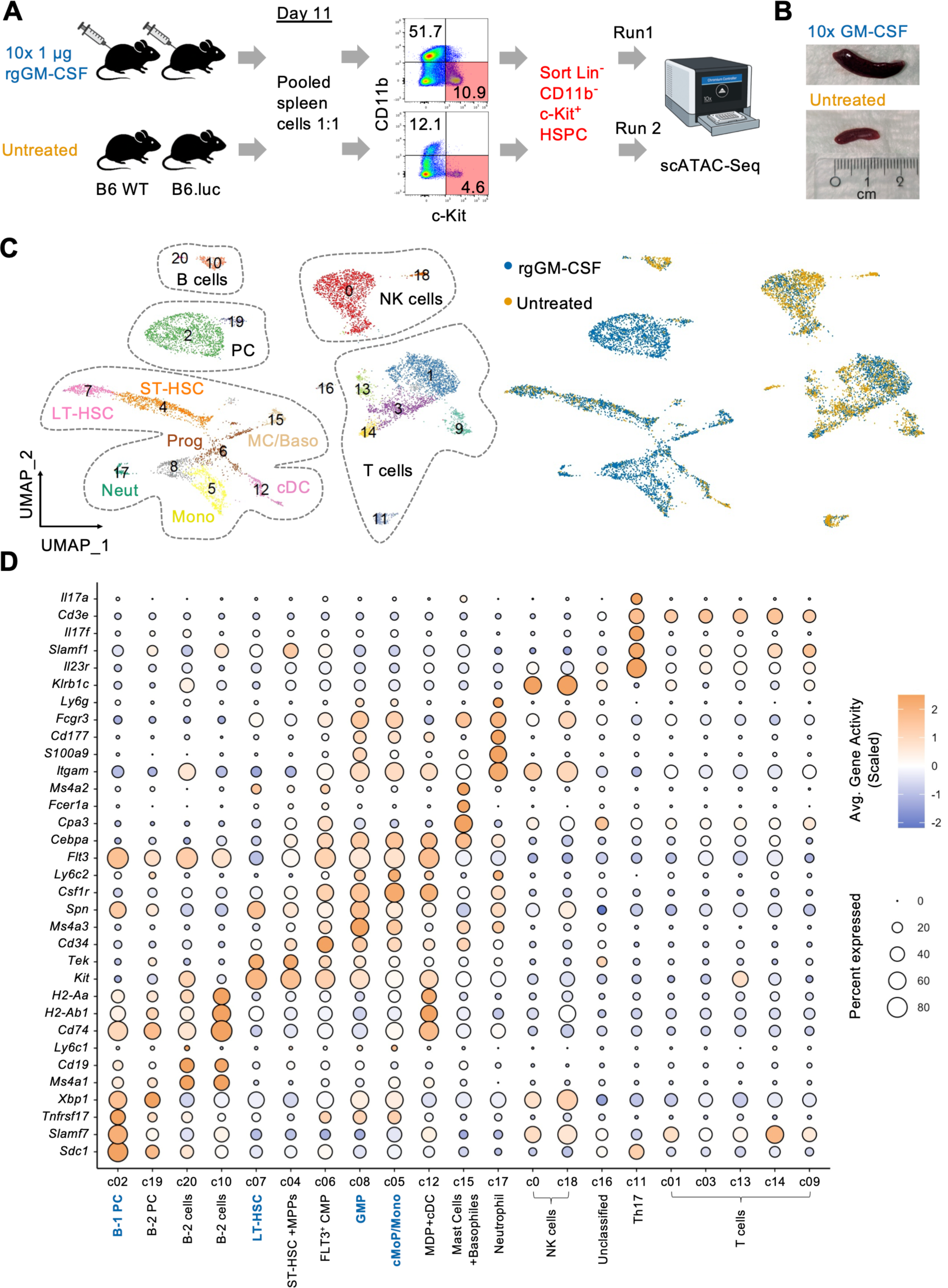
rgGM-CSF promotes splenic extramedullary hematopoiesis and B-1b plasma cell expansion in the spleen. (A) Experimental design for scATAC-seq. B6 albino and B6.lucefirase mice were injected intraperitoneally for 10 consecutive days with 1 µg rgGM-CSF (from supernatant from GM-CSF-producing cell line) or were left untreated. On d11, spleens were harvested, groups pooled 1:1, and Lin^-^ CD11b^-^ c-Kit^+^ hematopoietic stem and progenitor cells (HSPCs) isolated by FACS cell sorting, followed by scATAC-seq in two independent runs. (B) Representative images of spleens from the untreated or rgGM-CSF-treated mice at d11. (C) UMAP plot of the scATAC-seq data clustered on chromatin accessibility and annotated based on gene activity scores to infer cell identity (left UMAP), with the same embedding colored according to treatment condition (right UMAP). (D) Dot plots displaying scaled gene activity scores, and percentage of cells expressing lineage-defining markers across identified clusters.

Unexpectedly, in addition to the observed HSPCs and myeloid cell expansion, injections of rgGM-CSF further promoted a massive expansion of a B-1b PC population (Fig. 1C, cluster 2), characterized as *Spn⁺* (CD43) *Sdc1⁺* (CD138) *Slamf7⁺* (CD319) *Xbp1⁺* but lacking the mature B-cell markers *Cd19* and *Ms4a1* (CD20) (Fig. 1D). Flow cytometric analyses of peritoneal and splenic cells from PBS- and rgGM-CSF-injected mice further confirmed these findings (Fig. 2A, B). In the peritoneal cavity, rgGM-CSF IP injections reduced the frequencies of conventional B-2 cells (CD19^+^ B220⁺ CD138⁻ CD43⁻), B-1a cells (CD19^+^ B220^-^ CD138⁻ CD43⁺ CD5⁺), while inducing around 6-fold expansion of B-1b cells (CD19^+^ B220-CD138⁻ CD43⁺ CD5⁻), and a more than 10-fold expansion of B-1b PCs (CD138⁺ CD43⁺ CD5⁻), but not the B-1a PCs (CD138⁺ CD43⁺ CD5⁺). Systemically, in the spleen, rgGM-CSF led to a modest expansion of B-1b and B-1b PCs, whereas no differences were observed in B-2, B-1a or their corresponding PCs (Fig. 2B, C). Together, scATAC-seq and flow cytometric data showed rgGM-CSF-induced expansion of HSPCs, myeloid cells, B-1b cells and B-1b PCs.

**Fig. 2:**
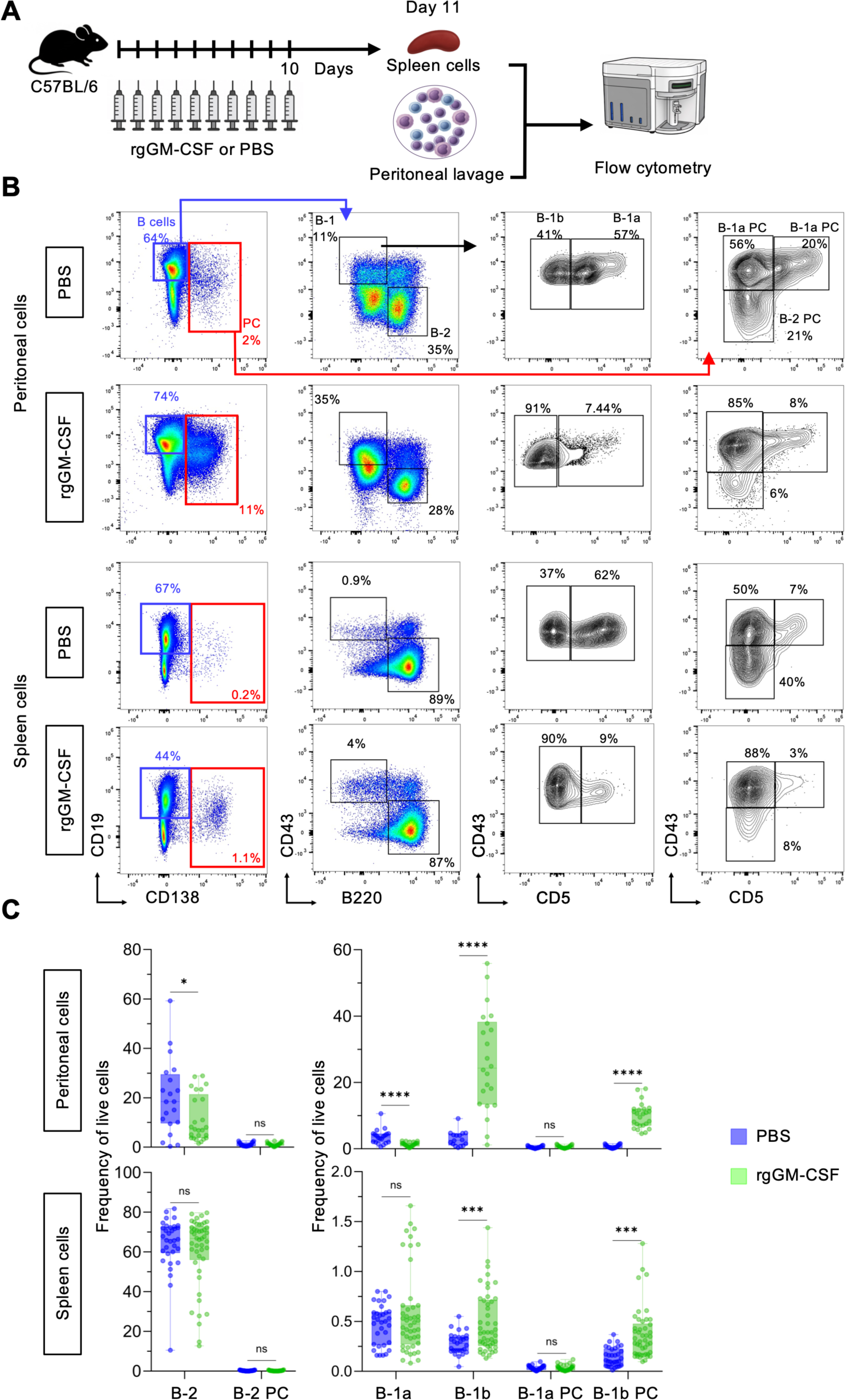
rgGM-CSF expands B-1b cells and B-1b plasma cells in the peritoneal cavity and spleen. (A) Experimental scheme showing the injection protocol, WT C57BL/6 mice were injected intraperitoneally for 10 consecutive days with 1µg rgGM-CSF or PBS. Peritoneal lavage and spleen cells were collected on day 11 for flow cytometric analysis. (B) Representative gating strategy of spleen and peritoneal cells for the identification of B-2, B-1a, B-1b, and their corresponding plasma cell populations pre-gated on single cells, live CD3^-^. (C) Quantification of frequencies of cell types indicated in (B) among live cells in the peritoneum and spleen. Statistical significance was determined by unpaired two-tailed *t*-test, n=6-12 biological replicates, with 2-3 technical replicates, 2-5 independent experiments. *p<0.05; ***p<0.001; ****p<0.0001, ns = not significant.

### rgGM-CSF-expanded LT-HSCs and MDSCs do not mediate protection from EAE

As we showed previously ^28^, injecting bulk BM rgGM-CSF cultures from WT mice as a source of M-MDSCs protected against EAE (Suppl. Fig. 1A, B). Since we identified iNOS-mediated nitric oxide secretion as the primary suppressive mechanism used by these M-MDSCs ^28^, we injected bulk cultures from *Nos2*^-/-^ mice. Surprisingly, the protection against EAE seemed independent of iNOS (Suppl. Fig. 1A, B). Additionally, transferring sorted neutrophilic or monocytic cells, the latter containing M-MDSCs, from the same cultures did not confer protection from EAE (Suppl. Fig. 1A, C, D), thus largely ruling out M-MDSCs as mediators of EAE protection. The scATAC-seq and flow cytometry data showed strong splenic expansion of HSPCs after 10 injections of rgGM-CSF (Fig 1C, Suppl. Fig. 1E, F). Therefore, GM-CSF-exposed HSPCs developing into suppressive myeloid cells could be candidates for EAE protection. However, neither adoptive transfer of sorted CD11b^-^ bulk progenitor cells from d3 BM rgGM-CSF cultures (Suppl. Fig. 1A, C, D) nor sorted splenic LT-HSCs from rgGM-CSF-injected mice (Suppl. Fig. 1G, H) proved protective, indicating that rgGM-CSF does not imprint HSPCs for suppression. In summary, these findings suggest that the sustained protective effect of rgGM-CSF is not mediated by HSPCs or M-MDSCs but may involve the previously unrecognized population of rgGM-CSF-expanded B-1b PCs.

### rgGM-CSF drives GM-CSF receptor-α expression and signalling pathways of B-1b PCs

To clarify whether B-1 cells can directly respond to GM-CSF we performed flow cytometric analysis of their GM-CSF receptor α-chain (GMCSFRα) expression. A significant fraction of the steady-state B-1a, B-1b, B-1a PCs and B-1b PCs, but not B-2 or B-2 PCs, expressed GMCSFRα, which was further increased after rgGM-CSF administration (Fig. 3A, B). To gain deeper insights into the GM-CSF signaling pathways in B-1b PCs, we performed GeneActivity analysis of the scATAC-seq data^36^. B-1b PCs exhibited marked upregulation of *Stat3, Stat5b, Mtor, Ei4ebp1, Pik3ca, and Irf1, Myd88* (Fig. 3C). Interestingly, the same GM-CSF signaling molecules we detected before in M-MDSCs ^28,45^. These data indicate that B-1 cells but not B-2 cells express the GMCSFRα and directly respond to rgGM-CSF.

**Fig. 3:**
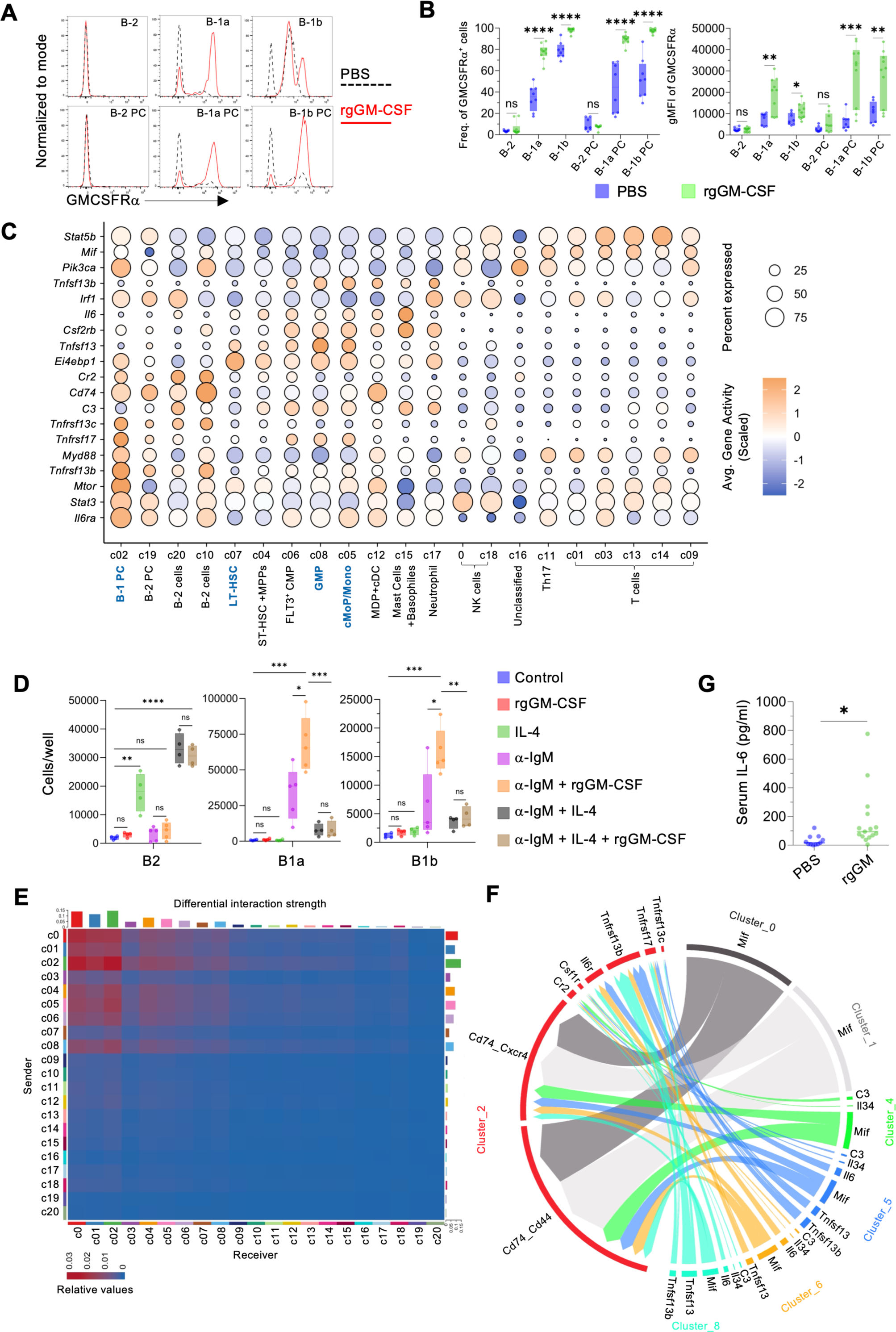
rgGM-CSF activates GM-CSF receptor signalling and a supportive myeloid niche for B-1b cells and B-1b plasma cells expansion. (A) Representative flow cytometric histograms showing expression of GM-CSFRα (stained intracellularly) on the indicated B-cell and plasma-cell subsets from PBS- or rgGM-CSF-treated mice following the 10-day injection protocol described in (Fig. 2 A). (B) Frequency of GM-CSFRα⁺ cells and geometric mean fluorescence intensity (gMFI) of GM-CSFRα among cell types indicated in (Fig. 2 B). (C) Dot plot of chromatin accessibility-derived gene activity scores for GM-CSF signaling pathway and B-cells-activating cytokines across scATAC-seq clusters shown in (Fig. 1C). (D) *In vitro* culture of splenic cells with anti-IgM, IL-4, and/or rgGM-CSF showing differential expansion of B-1 and B-2 cells. (E) Heatmap depicting CellChat-derived differential interaction strengths between scATAC-seq clusters shown in (Fig. 1 C). (F) Chord diagram illustrating predicted ligand-receptor interactions supporting B-1b plasma cell survival and activation. (G) Serum IL-6 concentrations measured by LEGENDplex™ on day 11 following the 10-day injection protocol outlined in (Fig. 2 A). For (B), n=8-10 biological replicates, 3-4 independent experiments. For (D) n=4-5 independent experiments. Statistical significance was determined by unpaired two-tailed *t*-test. *p<0.05; **p<0.01; ***p<0.001; ****p<0.0001, ns = not significant.

### rgGM-CSF requires co-stimulation and a supportive myeloid microenvironment to drive B-1 cell expansion

To uncover the mechanism underlying the rgGM-CSF-induced B-1b PC expansion observed *in vivo*, we cultured splenic cells *in vitro* with rgGM-CSF in the presence or absence of anti-B cell receptor antibodies (anti-IgM) and IL-4. Despite its striking *in vivo* effect, rgGM-CSF alone was unable to expand B-1 cells *in vitro* and required cooperation with anti-IgM to support the expansion of both B-1a and B-1b cells. Conversely, rgGM-CSF could not promote B-2 cell expansion, which was dependent on IL-4 and further enhanced by combination with anti-IgM (Fig. 3D).

Next, we examined the signals that may account for GM-CSF-mediated expansion of B-1b PCs *in vivo*. Their GeneActivity profile exhibited elevated gene activity scores for the BAFF receptor (BAFF-R, *Tnfrsf13c*), the APRIL receptors BCMA (*Tnfrsf17*) and TACI (*Tnfrsf13b*), the IL-6 receptor (*Il6ra*), and the complement receptor type 2 (*Cr2*) (Fig. 3C). Several GM-CSF-induced myeloid populations, including GMPs, cMoP/monocytes, and neutrophils (clusters 8, 5, and 17, respectively), showed coordinated gene upregulation of the corresponding ligands BAFF (*Tnfsf13b*), APRIL (*Tnfsf13*), C3 and IL-6 suggesting the presence of a supportive micro-environmental axis for B-1b PC expansion (Fig. 3C).

CellChat analysis ^38^ also confirmed that B-1b PCs (cluster 2) occupy a central position within an active intercellular communication network in the spleen. They received activating signals from multiple immune populations, including NK cells (cluster 0), T cells (cluster 1), ST-HSCs (cluster 4), cMoPs/monocytes (cluster 5), CMPs (cluster 6), and GMPs (cluster 8) (Fig. 3E). These populations produced ligands that paired with receptors on the B-1b PCs, such as macrophage migration inhibitory factor (MIF) with CD74, C3 with Cr2, IL-6 with IL-6R, and APRIL or BAFF with TACI or BAFF-R, together forming a composite signalling network able to sustain the expansion and activation of B-1b PCs *in vivo* (Fig. 3F). Mast cells and basophils (cluster 15) also showed elevated IL-6 expression (Fig. 3C), that is known to enhances PC viability and antibody secretion ^46^, which is in line with the detected increase of circulating serum IL-6 levels in rgGM-CSF-injected mice (Fig.3G).

### Albumin binding and glycosylation enhance the systemic bioactivity GM-CSF to drive B-1b PC expansion

To address the relatively weak *in vivo*-expansion of splenic B-1b PCs by injecting rgGM-CSF, compared to the strong effect observed with the in house GM-CSF preparation (GM-sup) used in the scATAC-seq experiment, we hypothesised that albumin present in the FCS of the GM-sup might have enhanced the systemic activity of GM-CSF. Since clinically approved human GM-CSF formulations are either not or only partially glycosylated, we also investigated whether glycosylation could influence B-1b cell expansion. To test both possibilities, we injected rngGM-CSF or rgGM-CSF with or without BSA for 10 d and monitored local (peritoneal) and systemic (splenic) B-cell responses by flow cytometry. Locally in the peritoneum, injections of rgGM-CSF alone were superior to enhance B-1b cell and B-1b PCs expansion than rngGM-CSF alone. BSA combinations with rngGM-CSF or rgGM-CSF expanded B-1b cells more effectively than the cytokines alone, while the frequencies of other cell types were reduced or remained stable (Fig. 4A). Systemically in the spleen, rngGM-CSF strikingly failed to induce the expansion of B-1b cells and B-1b PCs as observed with rgGM-CSF. This limitation of rngGM-CSF was compensated by co-injecting with BSA as a carrier protein, which slightly enhanced its systemic bioactivity and promoted B-1b cells and B-1b PCs expansion, although to a lesser extent than the rgGM-CSF/BSA combination (Fig. 4B). In addition, measurement of serum GM-CSF levels revealed that mice receiving rgGM-CSF ± BSA for 10 d exhibited significantly higher concentrations than those treated with rngGM-CSF ± BSA (Fig. 4C).

To exclude the possibility that BSA acted as a foreign antigen capable of eliciting B-1b PCs responses, we used normal mouse serum (NMS) as a source of mouse albumin. The NMS combination with rgGM-CSF also massively expanded B-1b cells and B-1b PCs in the peritoneum and spleen to levels comparable to those obtained with the BSA combination (Suppl. Fig. 2A and B), ruling out antigenic effects of BSA. Taken together, rgGM-CSF shows a higher systemic bioactivity than rngGM-CSF, correlating with its elevated serum levels and its capacity to expand splenic B-1b and B-1b PCs. Both bovine and murine albumin further increase the systemic activity of both glycoforms in the spleen, augmenting frequencies of B-1b cells and B-1b PCs.

**Fig. 4:**
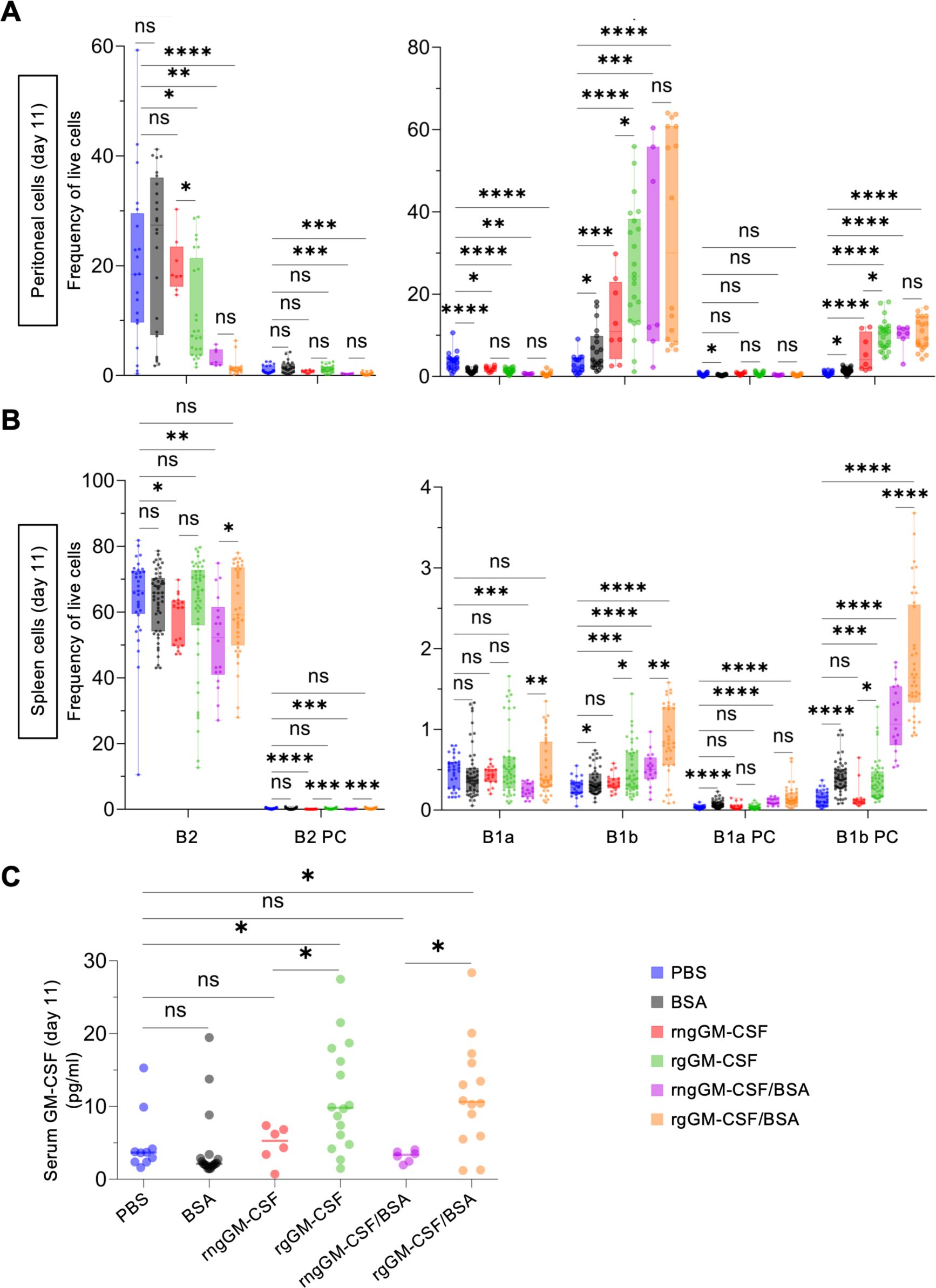
Glycosylation and albumin binding enhance systemic bioactivity of GM-CSF. (A, B) Frequencies of B-cells and plasma-cell subsets indicated in (Fig. 2 B) in the peritoneal cavity (A) and spleen (B) following 10 days of IP injections of 1 µg rgGM-CSF, rngGM-CSF, rgGM-CSF or rngGM-CSF in combination with 8 mg BSA, 8 mg BSA alone, or PBS. (C) Serum GM-CSF concentrations measured by LEGENDplex™ on day 11 following the 10-day injections described in (A, B). Statistical significance was determined by unpaired two-tailed *t*-test, n=3-12 biological replicates, with 2-3 technical replicates, 2-5 independent experiments. *p<0.05; **p<0.01; ***p<0.001; ****p<0.0001, ns = not significant.

### rgGM-CSF primes B-1b PCs for potent immunoregulatory function

To address the presumed suppressive function of the expanded B-1b PCs, Gene Ontology analysis of the top 200 upregulated genes in Cluster 2 was performed and revealed enrichment for immune-regulatory pathways (Fig. 5A). GeneActivity analysis for major immunoregulatory molecules ^47–50^ showed that rgGM-CSF injections elicited a strong induction of *Il10, Cd274* (PD-L1), *Ctla4, Tgfb1*, and *Arg1* selectively in B-1b PCs (cluster 2) (Fig. 5B). Flow cytometric analysis of peritoneal cells demonstrated that B-1a and B-1b cells and their corresponding PCs, but not B-2 cells upregulated the inhibitory receptors LAG3 and PD-L1 on their surface after rgGM-CSF injections (Fig. 5C-F). Consistent with the role of B-1 cells in natural IgM production for maintaining immune tolerance, rgGM-CSF-treated mice showed increased serum α-FN1, α-MDA and α-PC IgM titers at d11, which were maintained until d22. In contrast, rngGM-CSF did not or only minimally elevated their serum levels (Fig. 5G).

**Fig. 5.**
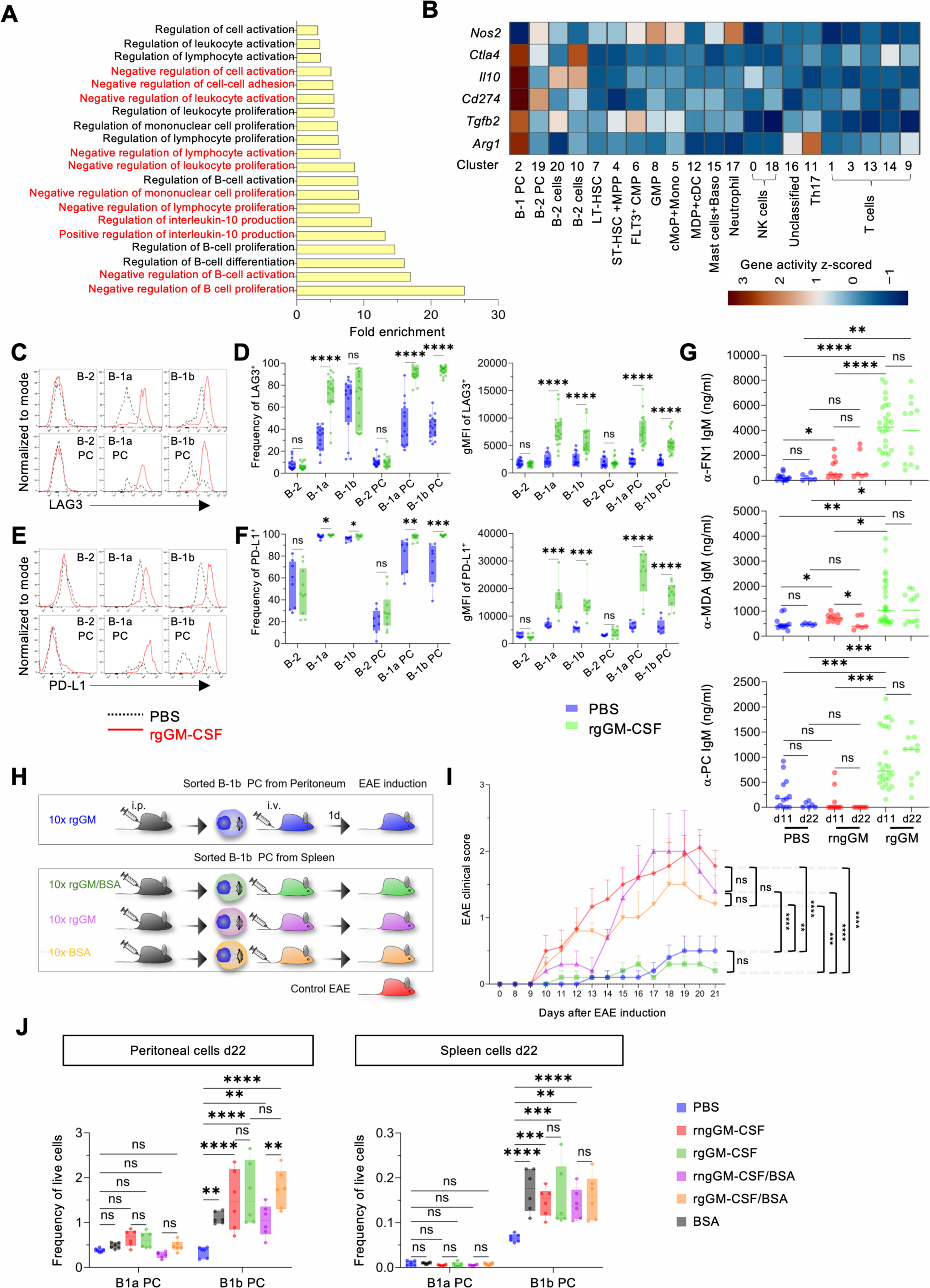
rgGM-CSF programs B-1b plasma cells for immunoregulatory function. (A) Gene ontology enrichment analysis of the top 200 upregulated genes obtained from chromatin accessibility-derived gene activity scores in the B-1b plasma cell cluster (Cluster 2) shown in (Fig. 1C). (B) Heatmap of gene activity z-scores for selected immunoregulatory molecules across scATAC-seq clusters shown in (Fig. 1C). (C) Representative flow cytometric histograms showing surface expression of LAG3 on the indicated B-cell and plasma-cell subsets from PBS- or rgGM-CSF-treated mice following the 10-day injection protocol described in (Fig. 2 A, B). (D) Frequency of LAG3⁺ cells and geometric mean fluorescence intensity (gMFI) of LAG3 among cell types indicated in (Fig. 2 B). (E) Representative flow cytometric histograms showing surface expression of PD-L1 on the indicated B-cell and plasma-cell subsets from PBS- or rgGM-CSF-treated mice following the 10-day injection protocol described in (Fig. 2 A, B). (F) The frequency of PD-L1⁺ cells and geometric mean fluorescence intensity (gMFI) of PD-L1 among cell types are indicated in (Fig. 2 B). (G) Serum levels of selected natural IgM antibodies specific for fibronectin (α-FN1), phosphorylcholine (α-PC), and malondialdehyde (α-MDA) were measured by ELISA at d11 and d22 following 10 days of daily injections of PBS, 1 µg rgGM-CSF, or rngGM-CSF. (H) Experimental design for adoptive transfer of 1x10^5^ MACS-sorted CD138⁺ B-1b plasma cells isolated from the peritoneal cavity or spleen of wild-type (WT) mice treated for 10 days with 1 µg rgGM-CSF, 8 mg BSA alone, or rgGM-CSF in combination with BSA, into recipient WT mice one day prior to EAE induction. (I) Clinical EAE scores following adoptive transfer as outlined in (H). (J) Frequencies of B-1 plasma cells in the peritoneum and spleen at d22 following 10 d of IP injections of 1µg rgGM-CSF, rngGM-CSF, rgGM-CSF or rngGM-CSF in combination with 8 mg BSA, 8 mg BSA alone, or PBS. For (D, F, G), statistical significance was determined by unpaired two-tailed *t*-test, n=6-14 biological replicates, 2-5 independent experiments. For (I) Data are shown as mean ± SEM. Statistical significance was determined by two-way ANOVA with Tukey’s multiple comparisons, n=5 biological replicates. For (J) Statistical significance was determined by two-way ANOVA with Tukey’s multiple comparisons, n=3 biological replicates with 2 technical replicates.*p<0.05; **p<0.01; ***p<0.001; ****p<0.0001, ns= not significant.

To directly assess their suppressive capability, MACS-sorted CD138⁺ B-1 PCs, consisting almost exclusively of B-1b PCs (Fig. 4A, B) from the peritoneum or spleen of the rgGM-CSF-injected mice were adoptively transferred into naive recipients before EAE induction, and compared with transfers of CD138⁺ B-1 PCs from spleens of mice injected with rgGM-CSF/BSA or BSA. The results showed that only peritoneal B-1 PCs from rgGM-CSF-injected mice and splenic B-1 PCs from rgGM-CSF/BSA-injected mice conferred prolonged protection from EAE, whereas B1 PCs isolated from the spleens of BSA- or rgGM-CSF-alone-treated mice were not protective (Fig. 5H, 5I). These findings indicate that local rgGM-CSF is sufficient to imprint suppressive activity in peritoneal bona fide B-1b PCs, whereas systemic delivery of rgGM-CSF requires co-administration of BSA. Serum IgM levels and EAE protection observed over 3 weeks suggested the persistence of B-1b PCs. At d22 the frequencies of both B-1 PCs were reduced compared with d11 but remained elevated relative to PBS-treated controls in peritoneum and spleen (Fig. 5J). These data suggest that rgGM-CSF induces an immunoregulatory program in B-1b PCs for at least 3 weeks.

### EAE protection by rgGM-CSF-imprinted B-1b PCs depends on their IL-10 production

To address the suppressive mechanism in the EAE protection and confirm the high IL-10 signature in the GO terms and GeneActivity, we next investigated IL-10 expression and its putative suppressive function in rgGM-CSF-induced B-1b PCs. Peritoneal and splenic cells of IL-10-β-lactamase reporter mice (ITIB) injected with rgGM-CSF, BSA or both were restimulated with PMA/Ionomycin and stained for IL-10 (Fig. 6A). Flow cytometric analysis showed elevated frequencies of IL-10^+^ B-1b PCs from mice injected with rgGM-CSF, which was further boosted in the spleen by the BSA combination (Fig. 6B, C). To determine whether B cell-derived IL-10 contributed to EAE protection, B6.CD19-Cre x IL10^fl/fl^ mice, which lack IL-10 expression in CD19⁺ B cells, and control B6.CD19-Cre mice were treated with rgGM-CSF. Their peritoneal CD138⁺ PCs were then MACS-purified and adoptively transferred into recipient mice before EAE induction (Fig. 6D). Transferred CD138⁺ PCs lacking IL-10 production failed to protect from EAE (Fig. 6E), indicating that IL-10 is a dominant immunoregulatory mechanism imprinted by rgGM-CSF in B-1b PCs.

**Fig. 6.**
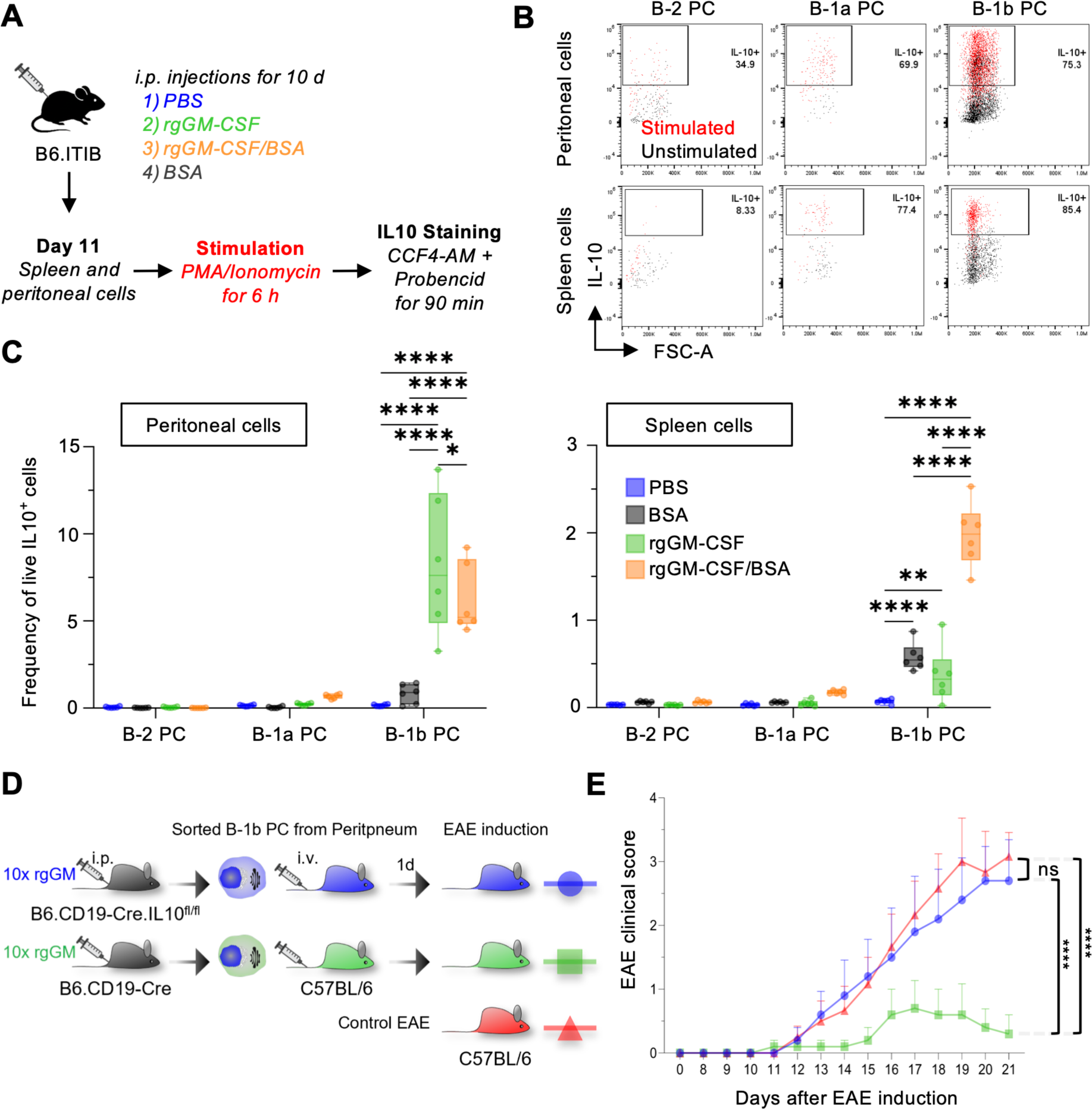
IL-10 production by rgGM-CSF-induced B-1b plasma cells mediates EAE protection. (A) Experimental scheme for 10-day IP injections of PBS, 1µg rgGM-CSF, 8 mg BSA alone or rgGM-CSF in combination with 8 mg BSA, into IL-10-β-lactamase reporter (ITIB) mice. At day 11, peritoneal and spleen cells were stimulated with PMA/Ionomycin for 6 h followed by IL-10 detection with CCF4-AM/Probencid for 90 minutes. (B) Representative flow cytometry plots showing IL-10 expression in peritoneal and spleen plasma cells from 10 days rgGM-CSF-treated miceafter PMA/Ionomycin stimulation and IL10 staining as outlined in A. (C) Quantification of IL-10⁺ plasma-cell frequencies in peritoneum and spleen after different injections, as indicated in A. (D) Experimental design for adoptive transfer of 1x10^5^ MACS-sorted CD138⁺ B-1b plasma cells isolated from the peritoneal cavity of B6.CD19-Cre.IL-10^fl/fl^ or B6.CD19-Cre mice after 10 d of daily IP injection with 1µg rgGM-CSF, into recipient WT mice one day prior to EAE induction. (E) Clinical EAE scores following adoptive transfer as outlined in (D). For (C), n=3 biological replicates with 2 technical repeats. For (E), n=5 biological replicates. Statistical significance was determined by two-way ANOVA with Tukey’s multiple comparisons. *p<0.05; **p<0.01; ***p<0.001; ****p<0.0001, ns= not significant.

### rgGM-CSF induces a distinct, transient expansion of iNOS^+^ M-MDSCs compared with rngGM-CSF

The scATAC-seq data and our previous results showed that rgGM-CSF administration induces M-MDSCs expansion in mice ^28^. To assess whether GM-CSF glycosylation had an influence on different myeloid cells (Fig. 7A), mice were treated with rgGM-CSF ± BSA, rngGM-CSF ± BSA, BSA or PBS. Injections of rgGM-CSF alone elicited a preferential increase in the frequencies of M-MDSCs and MoDCs at d11, whereas rngGM-CSF or rngGM-CSF/BSA showed a bias to elicit classical effectors of emergency myelopoiesis such as classical monocytes (Cl. Mono), macrophages, neutrophils, and Ly6C^lo^ neutrophil precursors (Fig. 7B). Following *ex vivo* stimulation at d11 with LPS/IFN-ψ, iNOS^+^ M-MDSC frequencies and iNOS expression from rgGM-CSF-injected mice were higher than those from mice injected with rngGM-CSF alone although this was partially compensated in the presence of BSA (Fig. 7C). Importantly, iNOS expression was no longer detected at d22 (Fig. 7E, F). In summary, rngGM-CSF rather promotes pro-inflammatory myeloid cells, while rgGM-CSF transiently supports the expansion and higher iNOS activity of M-MDSCs.

**Fig. 7:**
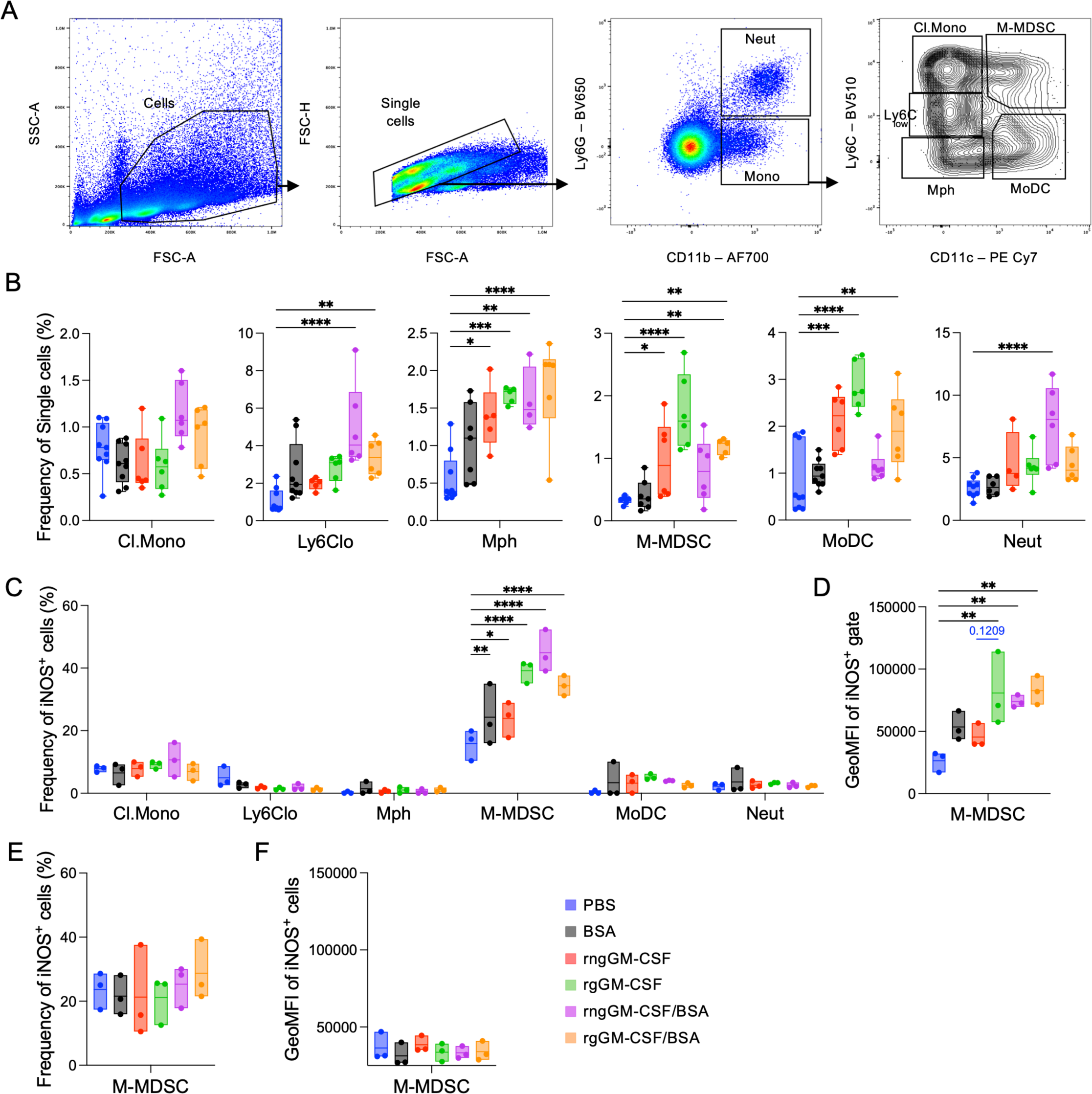
Preferential expansion of M-MDSC by rgGM-CSF, which shows the strongest iNOS expression in the spleen. C57BL/6 mice were treated shown in (Fig. 4A, B). Half of the mice from each group were sacrificed on d11 (B, C, D) and the other half on d22 (E, F). Splenocytes were harvested and analyzed by flow cytometry. (A) Gating strategy to identify neutrophilic (Neut) and monocytic (Mono) cells; Cells within the Mono gate were further divided into Cl.Mono, Ly6C^low^, macrophage (Mph), M-MDSCs, and MoDCs. (B) Frequencies of single cells of the gates indicated in (A). C, D: On d11, splenocytes were stimulated in vitro with LPS/IFN-γ for 24 h and then analyzed. Statistics showing the frequency of iNOS^+^ cells among the respective cell types (C) and the iNOS expression level per iNOS^+^ M-MDSCs (D). E, F: On d22, splenocytes were stimulated in vitro with LPS/IFN-γ for 24h and analyzed. Statistics showing the frequency of iNOS^+^ cells among total M-MDSCs (E) and the iNOS expression level per iNOS^+^ M-MDSCs (F). Statistics comparing PBS with all other groups. B, D, E, F: One-way ANOVA with Dunnett’s multiple comparisons test (black), unpaired t-test (blue), C: Two-way ANOVA with Tukey’s multiple comparisons, *p<0.05; **p<0.01; ***p<0.005; ****p<0.001. Only significant differences are indicated.

## Discussion

Although GM-CSF has long been recognized as a key regulator of myelopoiesis during inflammation, its opposing pro- and anti-inflammatory effects in mice and humans have remained largely unexplained and have generally been attributed to dose, application mode, anatomic location, and inflammatory context, rather than to molecular mechanisms ^2,51^. In this study, we found that i) GM-CSF glycosylation differentially controls the generation of pro- and anti-inflammatory cell types, ii) rgGM-CSF massively expands B-1b cells and B-1b PCs, and iii) the rgGM-CSF-induced B-1b PCs are functionally imprinted for immunosuppressive functions upregulating LAG3, PD-L1, and self-specific IgM and mediate suppression of EAE by IL-10. A similar Il-10^+^ LAG3^+^ CD138^hi^ PC phenotype, derived from several B cell subtypes, including B-1a, has been described as NRPCs that rapidly respond after *Salmonella* infection ^26^. However, whether GM-CSF played a role in this context has not been addressed.

It has been previously demonstrated that B-1a and B-1b plasmablasts respond to pathogens by secreting GM-CSF ^23,52,53^. It has been shown that autocrine GM-CSF enhances natural IgM secretion by B-1a cells and pathogen clearance ^23^. We found that all B-1 cells and B-1 PCs express the GM-CSFRα, which is further upregulated by GM-CSF. However, only B-1b cells and B-1b PCs selectively expand after rgGM-CSF stimulation, with increased activity of downstream signalling molecules STAT3, STAT5, PI3K, Irf1, 4E-BP1, and MyD88, as previously established for M-MDSCs ^28,54^. Single-cell ATAC-seq and CellChat ligand-receptor analyses further revealed that GM-CSF-expanded cells, such as GMPs, cMoPs, and monocytes, exhibited increased expression of factors implicated in B-cell proliferation, activation, and survival, including BAFF, APRIL, and complement components ^54–56^. Correspondingly, receptors for these ligands were expressed on B-1b PCs, consistent with coordinated intercellular signalling. In addition, we found that serum IL-6 levels mildly increased following GM-CSF injection, a finding also supported by CellChat analysis. IL-6 has been shown to be crucial for PC viability and antibody secretion ^46^. Together, these data indicate that GM-CSF exposure is associated with intercellular signalling networks that accompany B-1b cell differentiation.

Differences in pharmacokinetics and pharmacodynamics have been reported among the various glycoforms of GM-CSF. Injected rngGM-CSF peaked higher but was cleared more rapidly to baseline within 20 h, while the rgGM-CSF showed lower but persisted serum levels for 48 h ^57,58^, similar to what we detected here. In addition, fully glycosylated GM-CSF exhibits the weakest binding capacity to its receptor compared with partially glycosylated and non-glycosylated forms ^59^. Higher urinary leukotriene levels were reported after rngGM-CSF injection in humans than with the glycosylated form, underscoring its stronger pro-inflammatory activity ^57^. The systemic activity of GM-CSF is limited by its short half-life, but fusion to a carrier such as albumin can markedly enhance GM-CSF activity as shown in a tuberculosis model ^60^. We observed pronounced local B-1b and B-1b PC expansion in the peritoneal cavity for both glycoforms, whereas splenic responses were modest for rgGM-CSF and absent for rngGM-CSF. This constraint could be overcome by coadministration with BSA or NMS as a carrier protein leading to efficient splenic expansion.

GM-CSF is largely dispensable for steady-state myelopoiesis but is transiently induced during acute inflammation to initiate emergency myelopoiesis and expand innate effector cells ^2,6,61^. By contrast, sustained GM-CSF production in chronic inflammatory settings promotes ongoing myelopoiesis and the accumulation of immunosuppressive populations such as MDSCs [15,16]. Different dose-dependent functions of GM-CSF have been outlined extensively in tumor models and patients ^9^. Different dose-dependent signaling modes have been reported for the GMCSFR. Only high doses in a pro-inflammatory context allow the formation of dodecamer complexes and JAK2 signaling ^62,63^. Conversely, our previous findings support that low doses of GM-CSF generate tolerogenic or anti-inflammatory myeloid cells ^28,41,64^.

Our data suggest that these dose-related outcomes may result from differences in availability and glycosylation. The effects seen with rngGM-CSF, which is absorbed quickly and reaches high but transient serum levels, mirror the kinetics of emergency myelopoiesis, thereby mainly promoting the expansion of myeloid effector cells. In contrast, rgGM-CSF, which is absorbed more slowly and has lower peaks but maintains systemic exposure, favors differentiation towards immunoregulatory populations, including B-1b PCs and MDSCs. These patterns align with observations made with human GM-CSF preparations. The fully glycosylated regramostim appeared less effective at generating myeloid effector cells in humans compared to the fully glycosylated molgramostim and the partially glycosylated sargramostim ^8^ and consequently has never been approved for use in stem cell transplantation recipients, where monocytes and especially neutrophils are essential to fight opportunistic infections.

In conclusion, GM-CSF glycosylation controls the generation of pro- or anti-inflammatory cells in mice. While non-glycosylated GM-CSF acts pro-inflammatory, the fully glycosylated form supports the expansion of immunoregulatory B-1b PCs and M-MDSCs. These findings encourage re-evaluation of glycosylated regramostim for the induction of anti-inflammatory cell types and novel therapeutic strategies.

## Acknowledgement

We thank Marion Heuer and Mélanie Villard for expert technical assistance during the project and Moutaz Helal for help with the bioinformatics.

## Author Contributions

HA, AA, INE, LC, FI, ZL and EZ performed research and analyzed data; HA and FE performed the scATAC-seq bioinformatic analysis; HS, AB, NH, BP, EZ and BEC contributed valuable methods or mouse strains; HS, AB, BP, BEC TG, FE and MBL designed research; HA, AA and MBL wrote the paper.

## Supplementary Figures

**Supplementary Figure 1:**
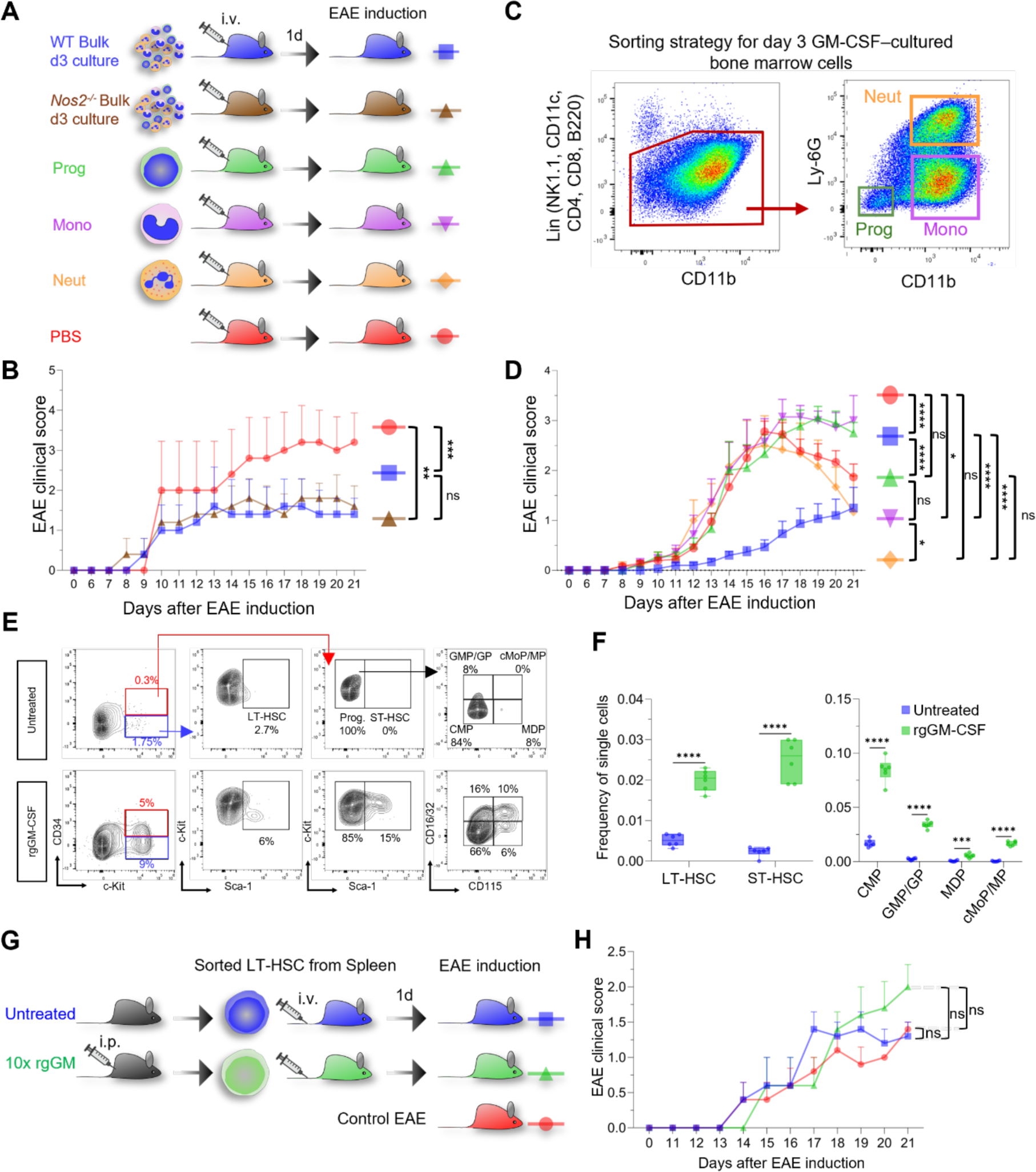
rgGM-CSF expands splenic HSPC but does not imprint long-term EAE protection. (A) Schematic of the EAE experiment involving adoptive IV transfer of bulk WT or *Nos2*⁻/⁻ BM cells from *in vitro* rgGM-CSF cultures for 3 d, sorted progenitors (Prog), monocytic cells (Mono) or neutrophilic cells (Neut) from the same culture as shown in (C), or PBS control, into recipient WT mice one day prior to EAE induction. (B, D) Clinical EAE scores across the indicated transfer conditions in A. (E) Flow cytometric gating strategy for identification of LT-HSC, ST-HSC, and myeloid progenitors in spleen cells at day 11 following 10 d of rgGM-CSF treatment or in untreated controls. (F) Quantification of frequencies of indicated splenic HSPCs populations shown in E. (G) Experimental design for adoptive transfer of 500 flow cytometry-sorted LT-HSC from untreated or 10 d rgGM-CSF-treated WT mice into recipient WT mice 1 day prior to EAE induction. (H) Clinical EAE scores following adoptive transfer as outlined in G. For (B, D and H) Data are shown as mean ± SEM, statistical significance was determined by two-way ANOVA with Tukey’s multiple comparisons, Mice/group used: B: n=5; D: n=9-18; H: n=5. For (F) statistical significance was determined by unpaired two-tailed *t*-test, n=6 biological replicates, 2 independent experiments, *p<0.05; **p<0.01; ***p<0.001; ****p<0.0001, ns = not significant.

**Supplementary Figure 2:**
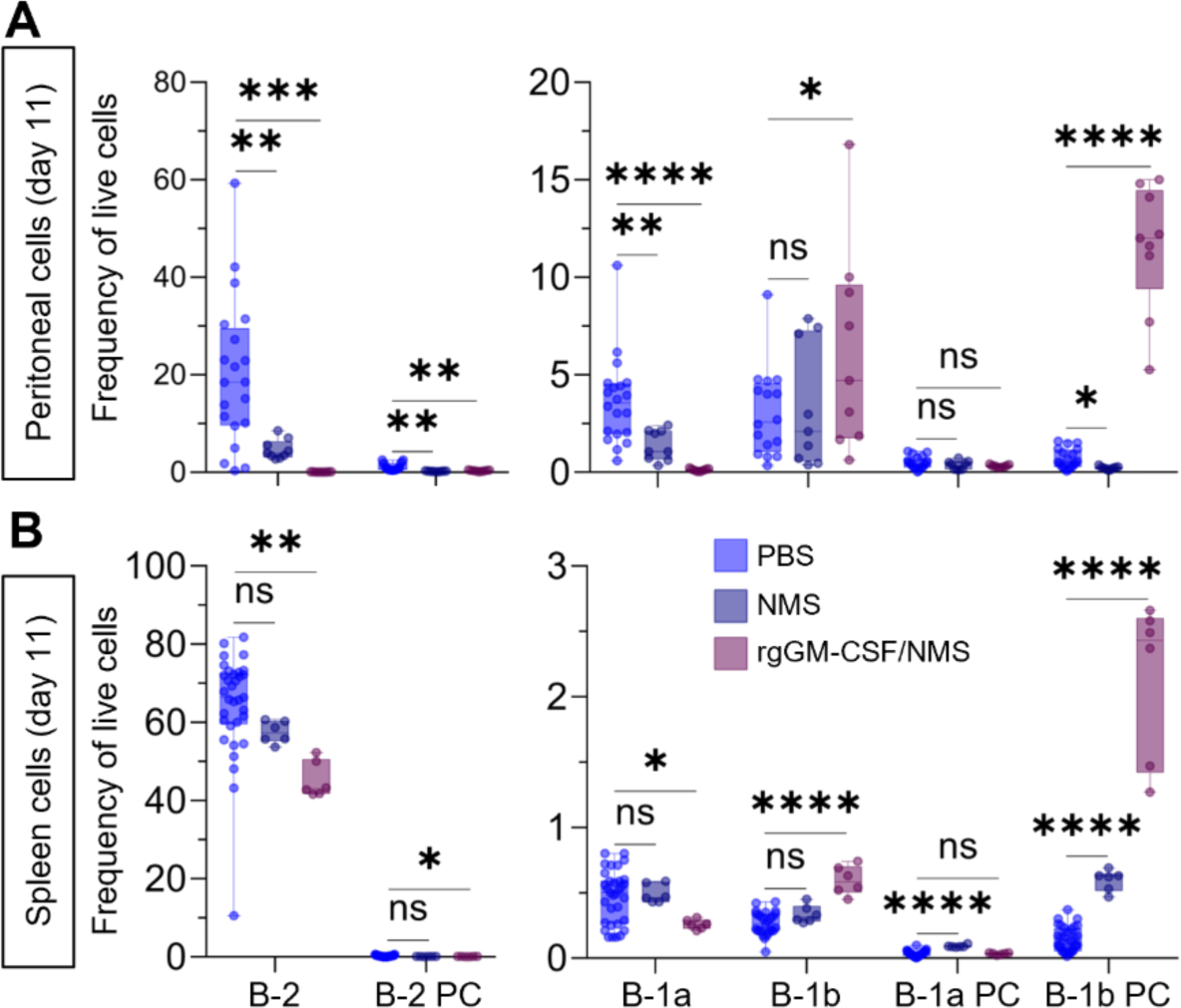
Mouse serum co-injection enhances the systemic rgGM-CSF-mediated expansion of B-1b cells and B-1b plasma cells in the spleen. (A, B) Frequencies of the cell populations indicated in (Fig. 2B) among live cells from the peritoneal cavity (A) and spleen (B) at day 11 following 10 days of daily intraperitoneal injections with PBS, 10% normal mouse serum (NMS) in PBS, or 10% NMS in combination with 1 µg rgGM-CSF. Statistical significance was determined by unpaired two-tailed *t*-test, n=3-12 biological replicates with 2 technical replicates. *p<0.05; ***p<0.001; ****p<0.0001, ns = not significant.

